# A Versatile Rapture (RAD-Capture) Platform for Genotyping Marine Turtles

**DOI:** 10.1101/450445

**Authors:** Lisa Komoroske, Michael Miller, Sean O’Rourke, Kelly R. Stewart, Michael P. Jensen, Peter H. Dutton

## Abstract

Advances in high-throughput sequencing (HTS) technologies coupled with increased interdisciplinary collaboration is rapidly expanding capacity in the scope and scale of wildlife genetic studies. While existing HTS methods can be directly applied to address some evolutionary and ecological questions, certain research goals necessitate tailoring methods to specific study organisms, such as high-throughput genotyping of the same loci that are comparable over large spatial and temporal scales. These needs are particularly common for studies of highly mobile species of conservation concern like marine turtles, where life history traits, limited financial resources and other constraints require affordable, adaptable methods for HTS genotyping to meet a variety of study goals. Here, we present a versatile marine turtle HTS targeted enrichment platform adapted from the recently developed Rapture (RAD-Capture) method specifically designed to meet these research needs. Our results demonstrate consistent enrichment of targeted regions throughout the genome and discovery of candidate variants in all species examined for use in various conservation genetics applications. Accurate species identification confirmed the ability of our platform to genotype over 1,000 multiplexed samples, and identified areas for future methodological improvement such as optimization for low initial concentration samples. Finally, analyses within green turtles supported the ability of this platform to identify informative SNPs for stock structure, population assignment and other applications over a broad geographic range of interest to management. This platform provides an additional tool for marine turtle genetic studies and broadens capacity for future large-scale initiatives such as collaborative global marine turtle genetic databases.

## Introduction

Marine turtles are migratory, long-lived megafauna of conservation concern, with populations of all species classified in high risk categories on the IUCN Red List of Threatened Species (IUCN 2017). The complex behaviors and life history traits marine turtles exhibit can make them highly susceptible to human impacts, while also posing challenges to understanding critical aspects of their biology required for their conservation (Wyneken *et al*. 2013). Over the past several decades, genetic approaches have provided key insight to important research questions in marine turtle biology and conservation, including natal homing to breeding grounds, connectivity between distant foraging grounds and nesting beaches, delineation of broad stocks and distinct population segments (DPS) for management (ESA 1973), and quantifying proportional impacts of fisheries across populations (reviewed in Jensen *et al*. 2013; Komoroske *et al*. 2017). Yet despite this progress, a diversity of unresolved research questions persist (Rees *et al*. 2016), many of which are well-suited to being addressed with emerging genetic and genomic approaches.

Genomic technological capabilities, especially high-throughput technologies (HTS), have rapidly expanded over the past decade to tackle a broader variety of questions in ecology and evolution (Ekblom & Galindo 2011; Ellegren 2014; Romiguier *et al*. 2014). Whole genome sequencing (WGS) and reduced representation approaches (such as targeted enrichment, transcriptome and restriction-site associated nuclear DNA sequencing; RNA-Seq and RAD-Seq, respectively) are becoming increasingly common with the continued decline in HTS costs and improvement of reference genome availability (Andrews *et al*. 2016; De Wit *et al*. 2015; Jones & Good 2016; Genome 10K 2009; Todd *et al*. 2016). However, resource development and applications in some taxa, especially many of conservation concern, have lagged behind others (Shafer *et al*. 2015; Garner *et al*. 2016). This is true for marine turtles and other non-mammalian vertebrates, highlighted by the fact that mammals comprise only 8% of the total number of vertebrate species, but represent over 70% of existing vertebrate genomes currently on *Ensembl* (Flicek *et al*. 2014). This has been in part due to limited resources and logistical constraints sampling animals with protected status and complex life histories, but also because these approaches are not compatible or cost effective with some of the highest priority research needs for these species. For example, WGS or reduced representation approaches that can be directly applied with little to no *a priori* genomic resources (RNA- and RAD-Seq) are well suited to address some research topics like phylogenomics and adaptive variation (Jarvis *et al*. 2014; Prince *et al*. 2017). However, other methods are needed for studies that necessitate background knowledge and tailoring approaches to yield informative variants (particularly single nucleotide polymorphism; SNPs) for specific study organisms and goals, such as research requiring cost-effective high-throughput genotyping data that are comparable over large spatial or temporal scales. This latter scenario is common in conservation research (Hunter *et al*. 2018) and monitoring of wide-ranging, long-lived species such as marine turtles, where samples often need to be compared across regions, continents and generations, such as fisheries bycatch DPS assignment and genetic capture-recapture studies (Komoroske *et al*. 2017; Shamblin *et al*. 2017; Stewart *et al*. 2016).

Several methods have recently emerged to meet these needs, including Genotyping-in-Thousands by sequencing (GT-Seq; Campbell *et al*. 2015), Rapture (RAD-Capture; Ali *et al*. 2016), and microhaplotypes (an adaptation of GT-Seq; Baetscher *et al*. 2017). Each of these approaches has demonstrated utility and strong potential for future broader application in conservation research under different study objectives and contexts. Marine turtle conservation researchers frequently encounter needs to genotype samples for different species, sample quantities, numbers of loci (e.g., for stock structure vs. relatedness studies), yet have limited time and financial resources to develop informative markers tailored to each study goal. Additionally, despite being one of the largest and most threatened vertebrate groups (Shaffer *et al*. 2015), there are currently limited reference genomes or transcriptomes for non-avian reptiles in general (but see Tzika *et al*. 2015; Shaffer *et al*. 2013; Wang *et al*. 2013), making it challenging to identify informative SNP loci *a priori* from existing genomic resources. Finally, researchers often deal with samples of varying tissue types, storage conditions, quality and quantity due to field, resource, and permitting and other limitations (e.g., samples from decomposing stranded animals, limited refrigeration in tropical study sites, and international CITES and shipping regulations). Thus, while no one approach provides an *a priori* solution to all of these research needs, we sought out to develop a robust, flexible platform that could be employed across a variety of research projects by adapting the Rapture method developed by Ali *et al*. (2016). In particular, we leveraged an existing molecular collection to test the utility of our approach with samples spanning the conditions frequently encountered in marine turtle research and combined initial RAD-Seq with Rapture target design to achieve this without *a priori* knowledge of good candidate regions. Here, we present our results and highlight the strengths, limitations, and future applications of this platform and general approach in marine turtle biology and conservation research.

## Materials and Methods

### Sample Selection, Processing and RAD-Sequencing

We selected 96 samples from the national Marine Mammal and Sea Turtle Research Collection (MMASTR) housed at NOAA Southwest Fisheries Science Center (La Jolla, CA) that collectively were representative of the genetic diversity among and within global leatherback populations. Samples were collected from 1988-2016, including nesting females, adult males, hatchlings (sex undetermined), as well as in-water foraging, stranded and bycaught animals of both sexes. Sample selection was weighted toward Pacific leatherbacks to contribute to a complementary project investigating fine-scale population structure in the Pacific. Tissue samples (skin, blood or muscle) were preserved in saturated salt when available, shipped, and stored in the NOAA-National Marine Fisheries Service MMASTR Collection at −20°C. Genomic DNA (gDNA) was isolated from sub-samples of tissue using one of the following standard extraction techniques: phenol/chloroform (Sambrook *et al*. 1989), sodium chloride (Miller *et al*. 1988), a modified DNeasy Qiagen extraction kit (Qiagen, Valencia, California), or Qiagen reagents on a Corbett CAS-1200 extraction robot (Corbett Robotics, San Francisco, California) or PerkinElmer JANUS robot (Waltham, MA). After extraction, gDNA was stored at −80°C until use in downstream analyses. All candidate samples were checked for DNA quantity and quality via Qubit Fluorometry (Thermo Fisher Scientific, Waltham, MA) and a 4200 TapeStation System (Agilent, Santa Clara, CA), respectively. Samples with adequate concentrations and the best quality (i.e., high molecular weight) were normalized and included in the final sample set for each location. Libraries were prepared following the updated RAD protocol as described in Ali *et al*. (2016) using *SbfI*-HF and NEBNext Ultra DNA Library Prep Kit for Illumina (New England Biolabs, Ipswich, MA) and sequenced at UC Davis Genomics Core Facility for paired-end 100 bp reads in 25% of a lane on an Illumina HiSeq 3000 instrument.

### RAD Data Analysis & Capture Target Design

We demultiplexed samples by assigning reads with complete matching barcodes (Ali *et al*. 2016) and assessed raw sequence data quality with FASTQC (Andrews 2010). The leatherback turtle genome has not yet been assembled, and the green turtle is the closest related species with reference genome. Although divergence of the *Dermochelidae - Cheloniidae* families is estimated at approximately 100 million years before present (Duchene *et al*. 2012), given the evidence for slower rates of DNA evolution among turtles relative to many other vertebrates (Avise *et al*. 1992) and the potential benefits of using a common reference genome relative to *de novo* assembly for our project goals, we aligned the leatherback RAD data to the green turtle genome (Wang *et al*. 2013) with the Burrows-Wheeler Aligner (BWA v0.7.5; Li & Durbin 2009) and evaluated mapping performance. We used *SAMtools* (v1.3; Li *et al*. 2009) to sort, filter for proper pairs and index alignments, remove PCR duplicates, and calculate summary statistics. After observing high mapping success (see results), we proceeded using these alignments to identify candidate SNPs and cross-species Rapture target loci. In brief, we employed a *SAMtools* genotype likelihood model in the program *ANGSD* (Korneliussen *et al*. 2014; Nielsen *et al*. 2012) to infer major and minor alleles and minor allele frequencies (MAF) for sites with data for at least one individual, mapping quality score ≥10 and base quality score ≥20. Specifically, we inferred major and minor alleles and estimated MAF using genotype likelihoods with a fixed major allele and unknown minor allele (Kim *et al*. 2011), adapted with an expectation-maximization algorithm as implemented in *ANGSD.* We then identified good candidate regions for targeted enrichment as regions with consistent coverage (∼84 bp length), paired both up and downstream of an identified restriction site in a high proportion of total individuals (≥68% for all samples; ≥80% for Pacific leatherbacks only), and without any suspected polymorphisms within the restriction site or unknown nucleotide identity *(N)* in the reference sequence. Within regions that passed these criteria, we then randomly selected one of the paired regions (i.e., either up- or downstream of the restriction site) and created candidate lists for two target types: (1) potential candidate SNP loci (MAF ≥0.1≤0.4, allowing only one variable site within 150bp from the restriction site; preferentially including those with a SNP within the first 84bp), and (2) no additional filters, to serve as a random locus set for unbiased genome representation within and across marine turtle species. We used corresponding sequences from the green turtle genome to design a custom MYBaits in-solution DNA target enrichment kit set (120bp baits, Arbor Biosciences, formerly MYcroarray Inc., Ann Arbor, MI) with ∼1000 targets for each of the two categories (2007 targets total) according to manufacturer protocols and quality control filters (e.g., probe compatibility, repeat masking, and melting temperature filters) with minor modifications to address initial failure of higher GC content baits (see below and Appendix S1 for details).

### Rapture Sample Selection, Library Preparation & Sequencing

We selected DNA samples from the MMASTR collection encompassing a cross section of covariates to examine the versatility of this method for the varied conditions frequently encountered in our studies (e.g., sample location, sex, life stage, collection method, tissue type, DNA concentration, DNA quality and collection year; 1342 samples total). In particular, we included samples with detectable concentrations at or below 5 ng/ul, which are frequently encountered in minimally invasive sampling of sensitive wildlife species, but below typical recommended concentrations for many reduced representation genome protocols. Although sample selection was again weighted toward leatherbacks for a complementary study, samples from six of the seven extant sea turtle species were included to evaluate target enrichment success across species and geographic regions, as well as green turtle samples representative of all currently defined global distinct population segments (DPS; Seminoff *et al*. 2015) to confirm the consistency of these genome-wide markers with established management delineations. We prepared RAD libraries as described above (Ali *et al*. 2016; 16 libraries total), with the modification of including samples with initial gDNA concentrations across the range frequently obtained from wild marine turtle samples (i.e., not selecting higher concentration samples only). A total gDNA of 50 ng was targeted as starting material for each library across all samples with a maximum input volume of 10 ul (i.e., samples with initial concentrations < 5 ng/ul had lower starting input). We quantified and normalized libraries, followed by targeted enrichment following manufacturer’s protocols, with the exception of doubling the capture reaction to include all RAD libraries (i.e., ∼1/8 capture reaction per RAD library). During amplification steps in RAD library and capture enrichment protocols, we estimated the minimum number of PCR cycles required for each library to minimize PCR clones.

The library enrichment process described above was conducted in two replicate trials after results from the first trial indicated a strong effect of GC bait content on enrichment success (Figure S1). After confirming with the manufacturer that our probe design m*et al*l quality control standards, a new, exact replicate MyBaits kit was synthesized. Library enrichment was repeated on the same RAD libraries with the new kit for Trial 2, along with minor amendments recommended by MYcroarray, Inc. to the original manufacturer protocol. For both trials, enriched libraries were combined and sequenced at the UC Davis Genomics Core Facility on an Illumina HiSeq 3000 instrument in a full lane (Trial 1: paired-end 100-bp reads, Trial 2: paired-end 150-bp reads). Here, except where specified, we focus on results from analyses of Trial 2 data only. However, we include a semi-quantitative comparison between the two trials with regards to on-target coverage to emphasize the importance of these technical details to inform effective MYBait design and application in future projects.

### Rapture Data Quality Assessment & Analyses

We demultiplexed samples as described above and assessed assignment error by quantifying the absolute and proportional number of raw reads (1) assigned to unused Illumina indexes or blanks (i.e., staggered wells without DNA within each plate/library) or (2) had barcodes on both forward and reverse reads. We assessed sequence data quality with *FASTQC* and *MultiQC* (Andrews 2010; Ewels *et al*. 2016), and calculated summary statistics in *R* (R Core Team 2016) to examine depth and evenness of coverage across predictor factors (e.g., library, species, tissue type, input concentration, sample location, and collection year). We used *BWA* and *SAMtools* as described above to map sequences and filter alignments. We qualitatively examined mapping quality using the *Integrative Genomics Viewer* (IGV; Robinson *et al*. 2011) and quantitatively assessed by locus and sample coverage at a representative position within target regions (relative position 20) with *Bedtools* (Quinlan & Hall 2010) and *R*. We combined information from raw read distributions and target loci coverage to establish quality (success/failure) thresholds, and only samples that passed these thresholds were included in subsequent data analyses. To quantify rates of on-target capture, we mapped forward reads to a reference of target loci only using the same pipeline described above with the exception of omitting PCR duplicate removal.

To examine and compare the success of our approach to generate SNPs within and across species and populations informative for various genotyping applications, we conducted SNP discovery, inferred major and minor alleles, and estimated allele frequencies for variable sites using *ANGSD* (Korneliussen *et al*. 2014; Nielsen *et al*. 2012) on a series of sample sets: (1) all turtle samples, (2) hardshell (Cheloniid spp.) turtles only, (3) green turtles only, (4) all leatherback samples, and (5) a representative leatherback population. For each sample set, we employed a genotype likelihood model and applied quality filters similar to RAD data as described above, additionally only including samples that passed initial QC thresholds and alignments that were proper pairs and uniquely mapped. Polymorphic sites were identified and retained in downstream analyses only if there were data for at least 50% of individuals within the group being tested, MAF ≥0.05, and p-value of being variable ≤1e-6. To examine relationships of coverage and predictor variables with genotyping success at multiple stringency levels, we estimated genotype posterior probabilities for a set of *a priori* candidate SNP positions (identified in RAD analysis described above) using an allele-frequency based prior and called genotypes with threshold cutoffs of 80, 90, and 95%.

### Species Confirmation & Population Structure Analyses

To validate our highly multiplexed approach, we first confirmed species identification with principal components analyses (PCA) by generating a covariance matrix without calling genotypes using the ngsCovar function in *ngsTools* (Fumagalli *et al*. 2014; Fumagalli *et al*. 2013) on all hardshell turtles, including a small sample set of suspected hybrids (based on morphological characteristics). To reduce influence of variance in depth of coverage between samples, we used *SAMtools* to randomly subsample alignments at multiple thresholds to balance information and sample retention in subsequent analyses (Ali *et al*. 2016). These analyses were also repeated including only less represented groups in the total hardshell dataset (i.e., loggerhead, olive ridley and Kemp’s ridley), where the higher proportion of green turtle samples could obstruct distinguishing variation. We also estimated admixture proportions of individuals using a maximum-likelihood-based clustering algorithm with the program *NGSAdmix* (Skotte *et al*. 2013) and genetic distances for a representative subset of samples across species and geographic regions using *ngsDist* (branch support based on bootstrapping 1000 replicates with 500 SNP blocks; Vieira *et al*. 2016) and plotted as a tree with *FastME* (BME iterative taxon addition method with NNI tree refinement; Lefort *et al*. 2015) and the R packages *phanhorn* (Schliep 2011) and *ape* (Popescu *et al*. 2012).

Secondly, we included green turtle samples from nesting grounds over a geographic range of interest to management in order to explore how our platform would perform delineating population structure within species. Thus, our goal was to evaluate the utility of the identified SNPs with this preliminary dataset to discern if they were likely to be informative markers in future, larger-scale analyses of stock structure and population assignment. We employed methods described above for PCA, admixture and genetic distances, and also estimated allele frequency spectra using *ANGSD* and *realSFS* to calculate pairwise F_ST_ values. Although it is common to accompany F_ST_ estimation with permutation tests to assess significant differences among the *a priori* defined groups, such analyses would have limited confidence given the restricted group sample sizes in our exploratory dataset, and are more suitable for future stock structure studies employing these markers with robust sample sizes and comprehensive geographic coverage.

Finally, we also estimated allele frequency spectra to calculate genetic diversity statistics (Watterson’s estimator, θ_w_, based on number of segregating sites, and Tajima’s estimator, θ_π_ or π, based on pairwise differences between sequences) in *ANGSD* and *realSFS* among species (Korneliussen *et al*. 2014; Korneliussen *et al*. 2013; Tajima 1989; Watterson 1975). Unequal sample sizes, population structure and upstream filtering for SNPs can cause biases in nucleotide diversity estimations (Lozier 2014; Subramanian 2016; confirmed with subsampling simulations on this dataset), potentially creating issues in our dataset with variable sample sizes across populations with likely differing demographic histories and current status (e.g., recovering, declining, etc.). To address this, we included only the random set of targeted loci as described above with selected subsets of 4-6 QC passed individuals from representative populations from each species, and report results on semi-quantitative evaluation of descriptive statistics only. Thus, although inference from these metrics is constrained, we include them demonstrate the utility of this platform for research employing these metrics in robust sample sets within or across species.

## Results

### RAD-Sequencing & Rapture design

We recovered 95.7 million total raw sequences, and 89.0% of which were retained based on sample assignment criteria. *FASTQC* confirmed consistent high sequence quality across the library with no evidence of contamination. After removal of four failed samples (defined as <2% of average number of sequences assigned to sample), an average of 93.9% (±7.3% S.D.) of sequences mapped to the green turtle genome, an average of 51.2% (±4.1% S.D.) of which remained after filtering out PCR clones. These results of strong concordance supported the use the green turtle genome as a reference, so we proceeded using these alignments for further Rapture bait development. We identified a total of 7,282 RAD tags with paired regions that met initial filtering criteria. A total of 1,379 of these candidate regions further met our SNP criteria (see methods) and were included in bait design, as well as 1,400 additional randomly selected regions from this list. From these 2,779 final candidates, we were able to design a custom MYBaits kit that met MYcroarray’s QC criteria with 2,007 targets for Rapture genotyping in marine turtles.

### Rapture data quality analysis

In Trial 2, we recovered 396 million total raw sequences, with only 0.38% of these sequences removed due to assignment to unused Illumina indexes or the presence of barcodes on both forward and reverse reads. *FASTQC* and *MultiQC* results confirmed high quality scores across and within libraries and no issues of contamination. Assignment of raw sequences to blanks dispersed across libraries was extremely low (average= 245, min/max=27/818). Based on sequence count distributions, we determined an initial sample failure/success threshold of 10,000 raw sequences, which 1127 samples passed (84%; hereafter referred to as ‘QC passed samples’). Read counts varied across library and samples, but we did not observe any clear patterns of success or failure between input factors, particularly among species or DNA input. Samples more recently collected and with higher DNA initial concentrations more consistently passed initial quality thresholds, but many low concentration and older samples did as well.

### Rapture target coverage and genotyping success

Samples exhibited very high percentages of mapping and on-target sequence capture, with Trial 2 having even higher on-target success than Trial 1 (Fig. 1A & S1; see methods and Appendix S1 for details). For Trial 2 data, mapped filtered (PCR clones removed) fragments for QC-passed samples were an average of 20.8% (±6.9% S.D.) of the total sequenced fragments per individual, and this was correlated with sample initial gDNA concentration (Fig. 1B). Average coverage per locus in filtered QC-passed samples was 26.6 (±10.1 S.D.; min/max=0.9/99.1; see Fig. S2 for coverage distributions). Samples generally reached ≥ 4x coverage across loci with approximately 50,000-75,000 filtered alignments (Fig. S3a). However, we identified samples that passed initial QC thresholds, but had lowered numbers of filtered alignments and few Rapture loci covered at ≥ 4x (Fig. S3b), prompting us to implement an additional filter of a minimum of 5,000 filtered alignments in further downstream analyses. Of these new QC-passed samples (1097 total), we were able to genotype over 50% of *a priori* identified SNPs in Rapture loci at all posterior probability thresholds tested (Fig. 2a). Genotyping capacity increased with depth of coverage but began reaching saturation at approximately 150,000 sequenced fragments per individual (depending on posterior probability threshold and sample). However, genotyping capacity was also clearly affected by the relative position of the SNP within the Rapture locus region (Fig. 2b), displaying a distinct break at approximately relative position 100, despite the use of longer 150bp paired-end sequencing.

**Figure 1.**
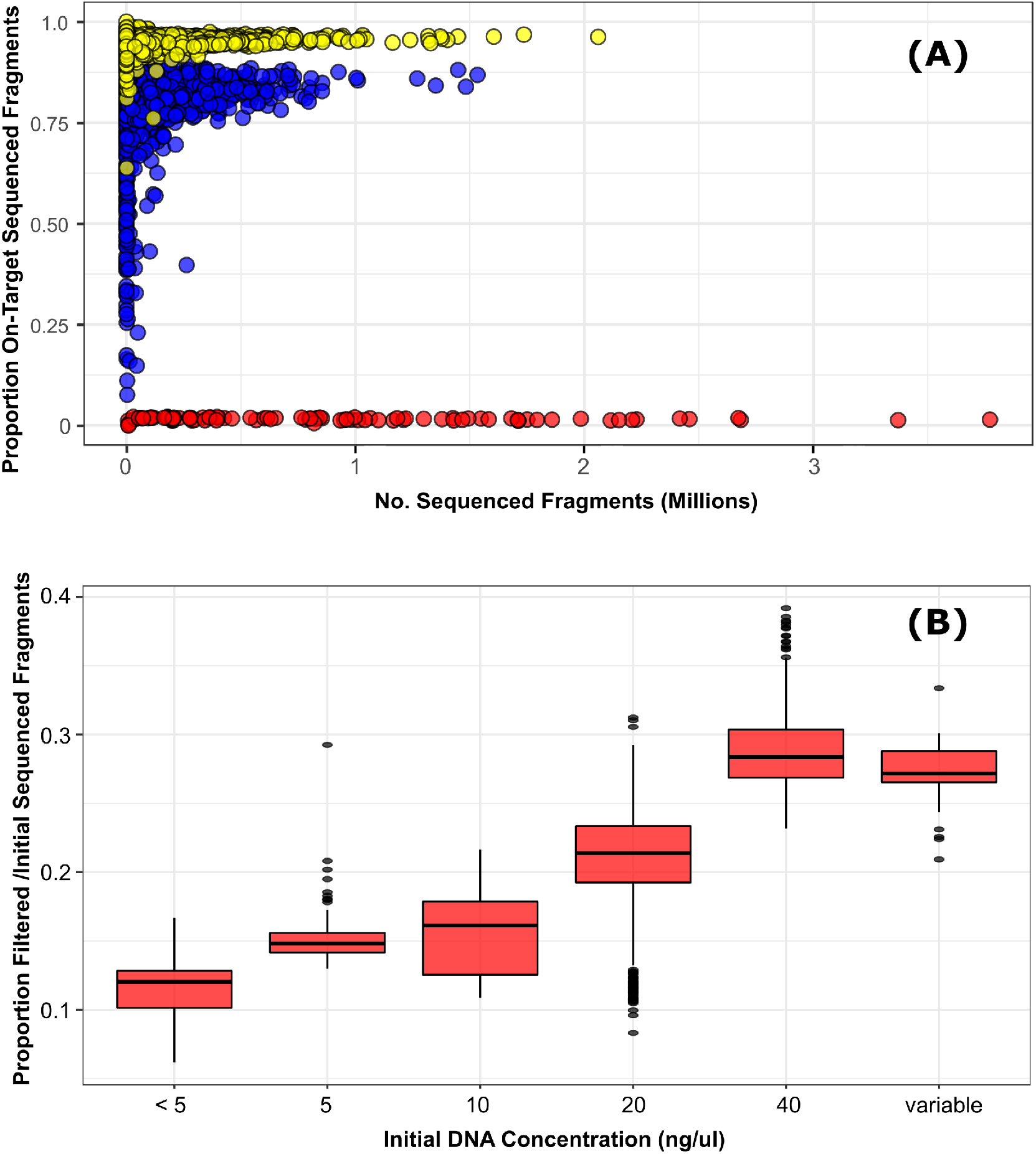
Panel (A) depicts the proportion of total sequenced fragments per individual that mapped to Rapture target loci from (1) initial RAD data (red circles), (2) Rapture data generated from original MYBaits protocol (Trial 1; blue circles), and (3) Rapture data generated from adapted MYBaits protocol (Trial 2; yellow circles). Note that one over-sequenced outlier with >7 million sequenced fragments was removed to improve visual interpretation. Panel (B) depicts the proportion of filtered mapped alignments/total sequenced fragments per individual for each category of initial DNA concentration (ng/ul).

**Figure 2.**
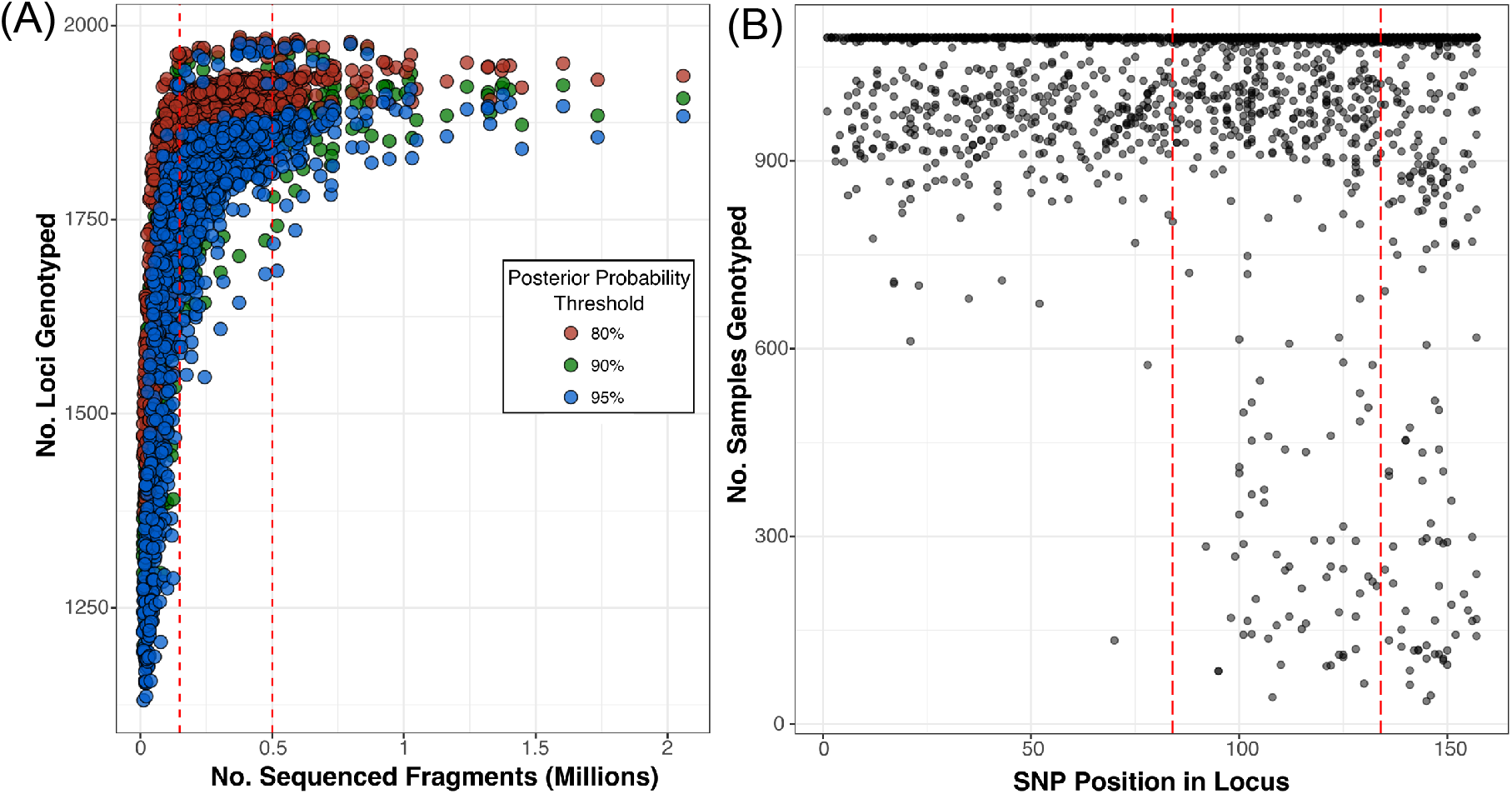
(A) Relationship between the number of sequenced fragments per individual and the number of *a priori* SNP loci genotyped, and (B) the relationship between the SNP relative position within a Rapture locus and the number of samples genotyped (visualized with 80% posterior probability threshold). Vertical lines added at relevant thresholds for visual interpretation (see text).

### Cross Species Capture Success & SNP discovery

We observed consistent success in coverage of Rapture loci across all species tested, confirming the broad utility of this approach for genotyping studies across marine turtle species. A reduction in the maximum loci covered regardless of total depth of coverage was observed in non-green hardshell turtle species (Fig. 3), indicating that a small percentage of selected targets in this particular enrichment set are not useful for other hardshell species, likely due to polymorphisms in *SbfI* restriction sites or other compatibility issues. Nevertheless, we identified ample candidate polymorphic SNPs suitable for within-species genotyping studies (Table 1). However, we emphasize that because SNP identification is inherently determined by analysis parameters and input sample composition, determining informative SNPs within Rapture target regions should be conducted using samples and filtering thresholds aligned with research goals to avoid ascertainment bias (e.g., demonstrated here by comparing SNP discovery results in all leatherback samples versus within one specific population; Table 1).

**Figure 3.**
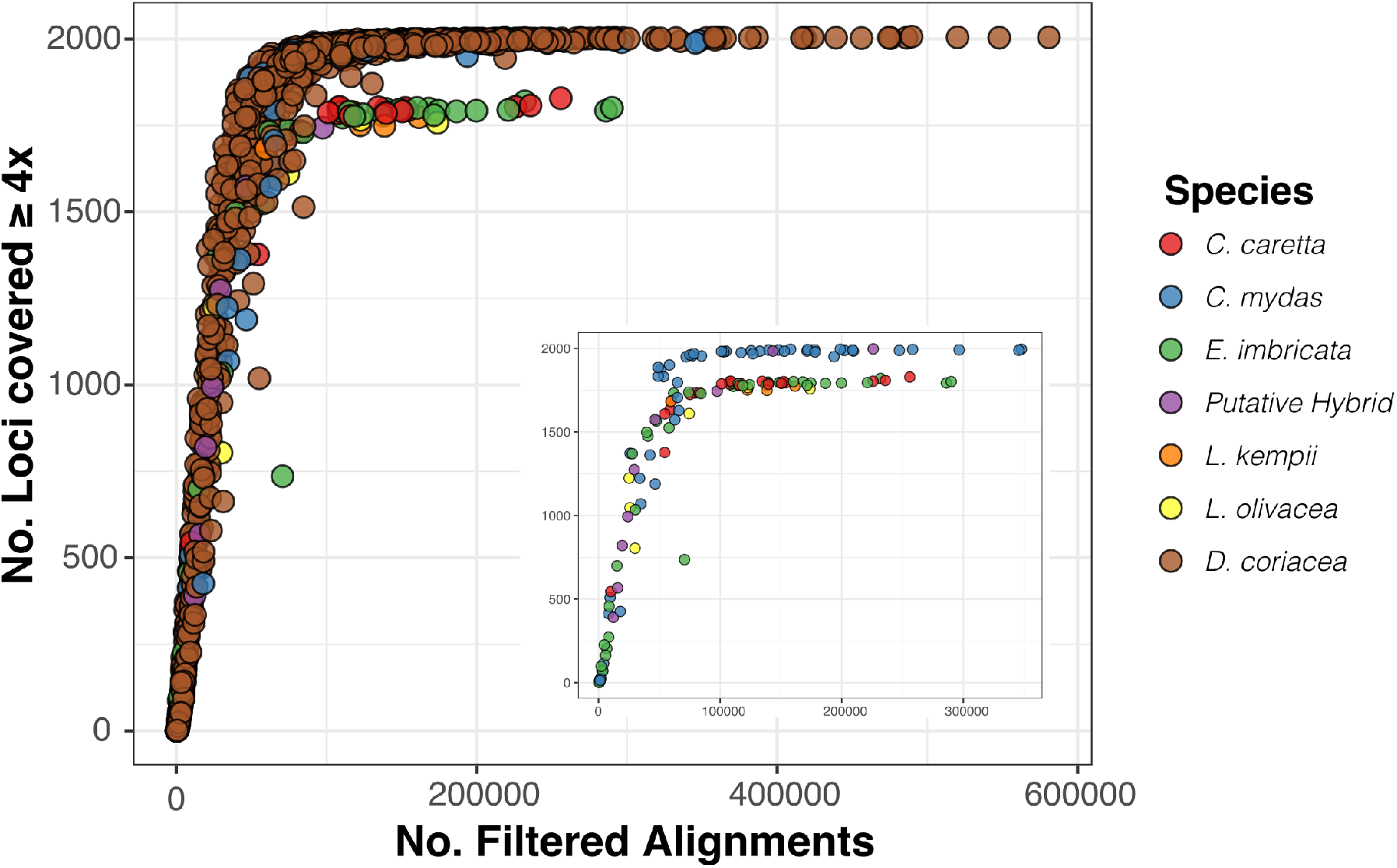
Number of Rapture loci covered ≥ 4x for all samples (one over-sequenced outlier with >1 million filtered alignments removed to improve visual interpretation). Inset depicts hardshell turtles to better visualize that only green turtles and green-hybrids attain coverage at all Rapture loci.

**Table 1.**
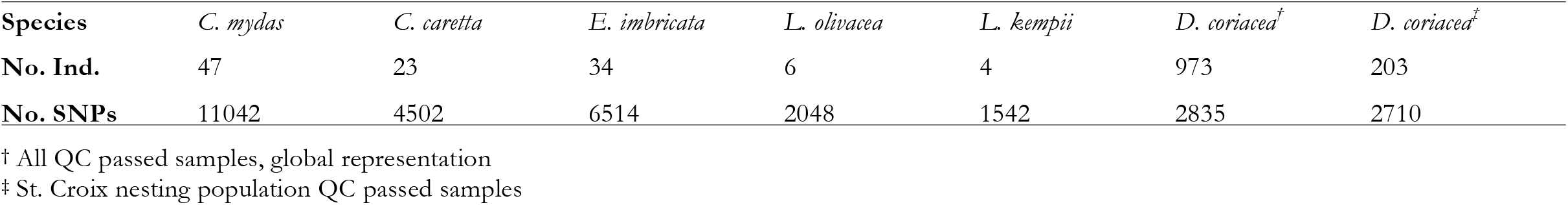
Initial SNP discovery per species with Rapture data for all QC passed samples (filters of MAF 0.05-0.4 and only sites with data for at least 50% individuals). Factors such as filtering thresholds, number of input samples, and source population of samples can affect identification of SNPs that are informative for different study goals.

### Species Confirmation and Green Turtle Population Structure

Individuals strongly separated by species as expected in the first two PC components for all hardshell species, with the exception of the two ridley species (Fig. 4a) that resolved in further PC axes in the combined analysis, as well as separate analyses omitting green and hawksbill turtle samples (Fig. 4b). Clear species separation was similarly observed in admixture proportion results, but with even more pronounced effects of the unbalanced sample groups when all hardshell samples were included (i.e., strong breaks in population structure within green turtles began to emerge before the separation of the ridley species; Fig. 4c,d). Estimated genetic distances among species were largest as expected between leatherbacks and hardshell turtles, followed by green turtles relative to other hardshell species (loggerhead, hawksbill, Kemp’s ridley, and olive ridley; Fig. S4). Several hybrids were identified, including three green-loggerhead hybrids and one green-hawksbill hybrid, however for several other suspected hybrids both PCA and admixture proportion results support only genetic contributions from olive ridley.

**Figure 4.**
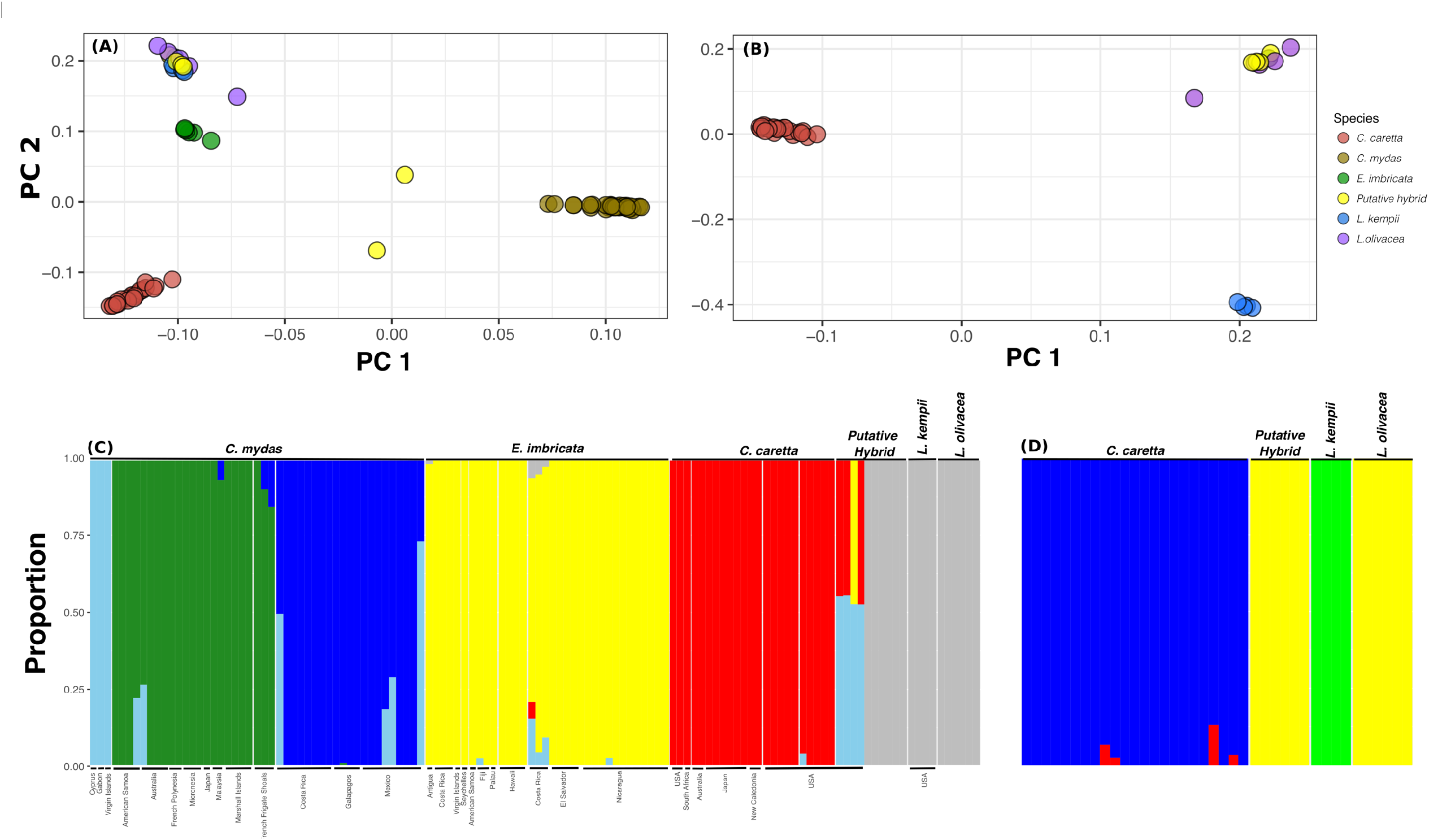
Species confirmation in hardshell turtles using principal components analyses (panels A and B) and admixture proportions (panels C and D). Panels (A) and (C) include all hardshell samples, while (B) and (D) include only of subsets of smaller groups, demonstrating how delineations among closer-related groups with smaller sample sizes can be masked in larger, disproportionate datasets. Only unresolved hybrids from the complete data set depicted in Panels A and C are included in Panels B and D.

In green turtles, pairwise F_st_ values, genetic distances and PCA discerned strong breaks in population structure between major ocean regions aligned with previous studies based on mtDNA and microsatellites and green turtle DPS designations (Jensen *et al*. in press; Seminoff *et al*. 2015; Figs. 5 & S5; Table S1). Tree topology branch support of genetic distances as well as F_st_ values were higher in the Atlantic compared to the Pacific Ocean. In the western Pacific, PCA clustering of samples by location for several groups are congruent with potential finer-scale population structure (Fig. S5b), further supporting the utility of these SNP markers for future stock structure and population assignment studies.

**Figure 5.**
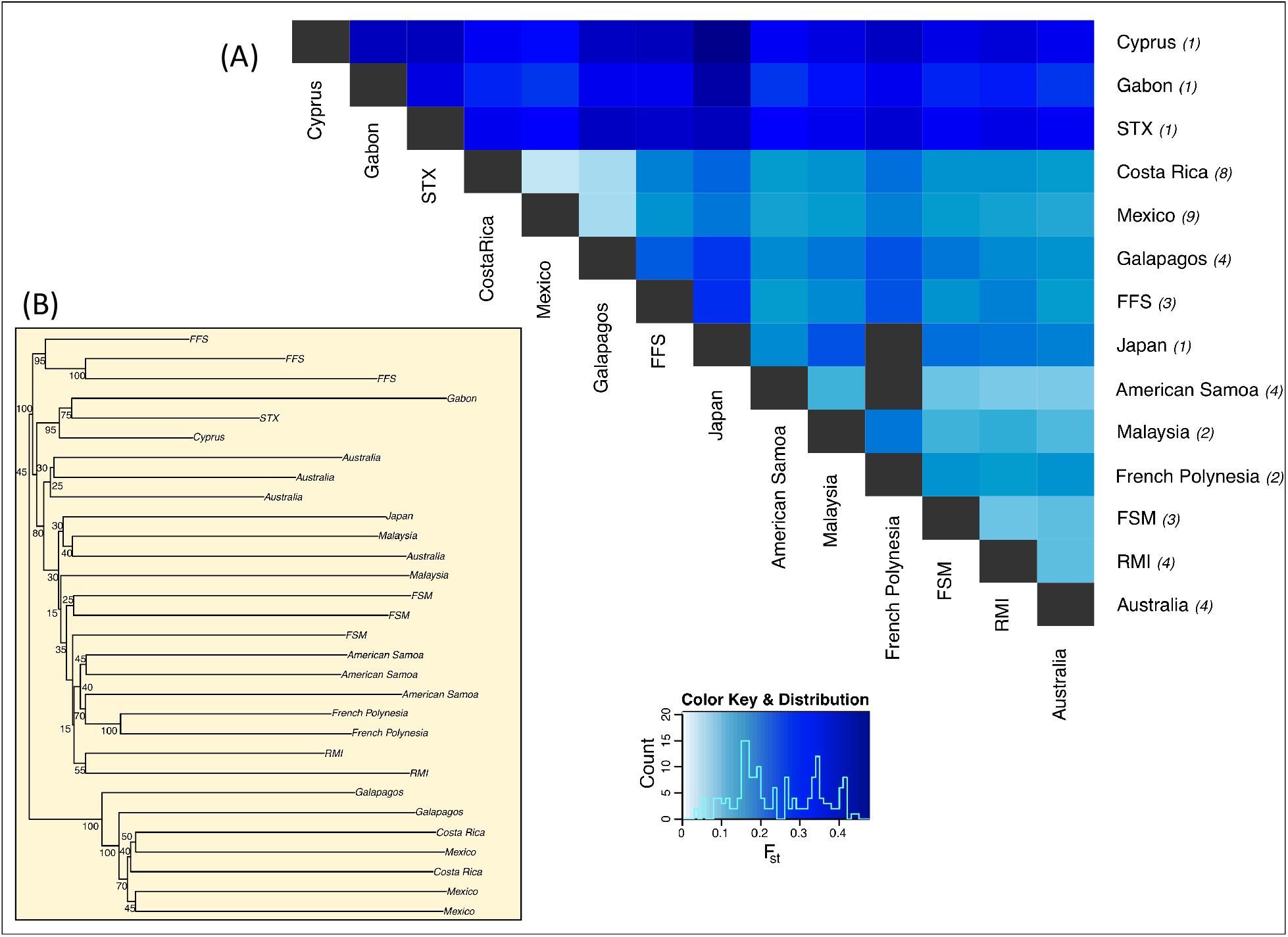
(A) Pairwise F_st_ values between green turtle nesting regions (sample sizes listed in italicized parentheses; black boxes indicates values could not be reliably calculated due to low sample size and sequencing coverage). (B) *FastME* tree of a representative subset of green turtle samples with topology and relative branch length based on genetic distances estimated in *ngsDist.* Branch support based on bootstrapping (1000 replicates, blocks of 500 SNPs). Abbreviations: STX=St. Croix, FFS=French Frigate Shoals, RMI= Republic of the Marshall Islands, FSM= Federated States of Micronesia.

### Genetic Diversity Estimates

Patterns within groups were consistent between θ_w_ and π, and within species, with the exception of Costa Rica hawksbills that had substantially higher values for both metrics (Fig. 6). Generally, green turtles exhibited the highest nucleotide diversity, while leatherbacks displayed the lowest. In particular, all four groups of Pacific leatherbacks had lower levels of variation relative to the Atlantic population included (Brazil).

**Figure 6.**
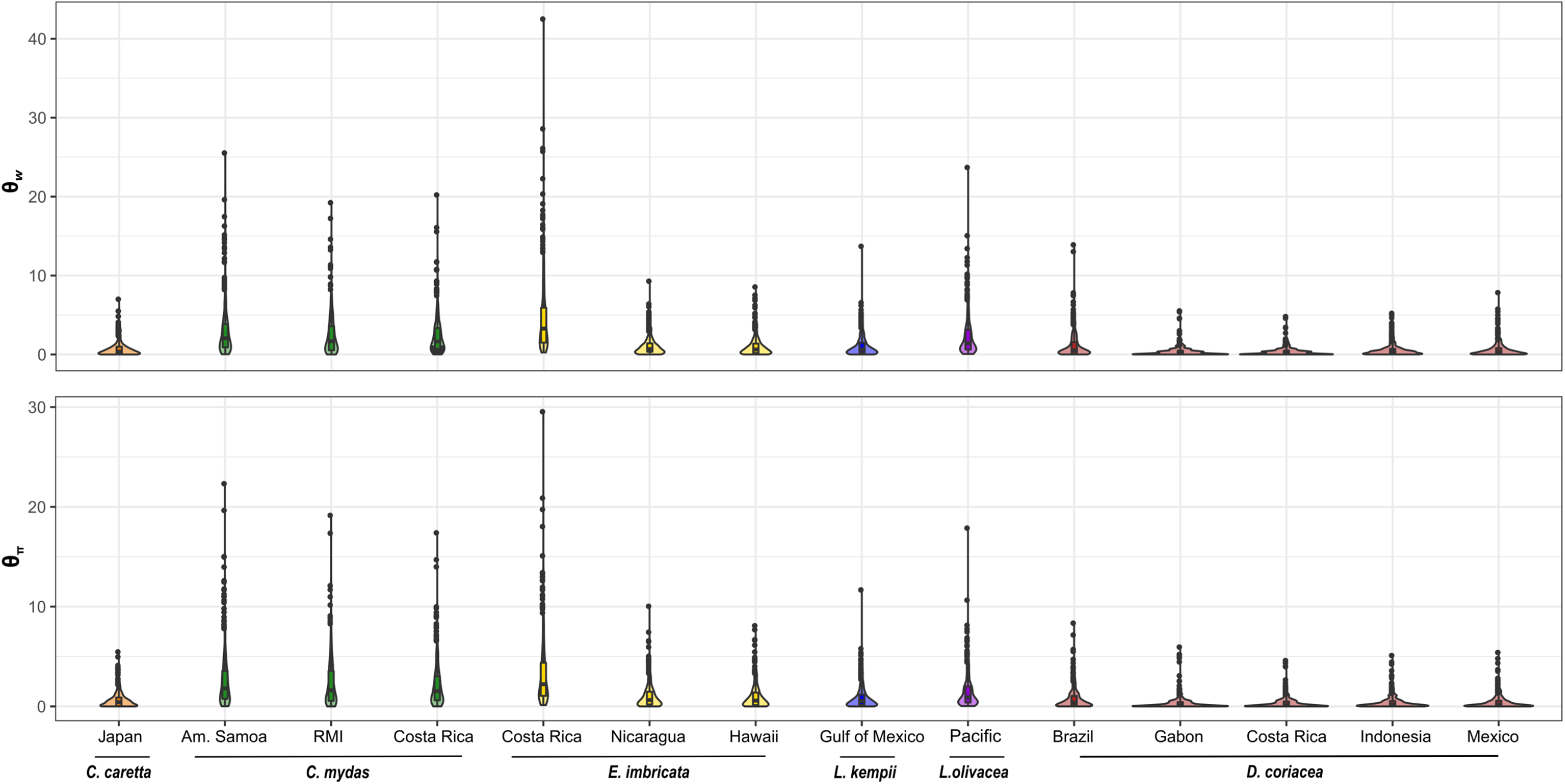
Genetic diversity estimates (top: Watterson’s estimator θ_w_; bottom: Tajima’s estimator θ_π_) in representative groups for each species. Locations listed indicate nesting population with the exception of *L. olivacea* for which only bycatch samples with unknown nesting origin were available.

## Discussion

Technological advances combined with increased interdisciplinary collaboration has rapidly expanded both the scope and scale of genetic studies over the past decade, yet for many species of conservation concern such as marine turtles, the realized potential of these advances is only just beginning (Garner *et al*. 2016; Komoroske *et al*. 2017; Shafer *et al*. 2015). This is in part because life history traits and protected status of these taxa can create unique research challenges, but also because the resources required for method development (which often needed to be repeated to generate informative markers tailored to each species and study goal) often has made it infeasible for conservation researchers. Our results demonstrate that the adaptation of the Rapture method developed by Ali *et al*. (2016) provides a flexible platform for marine turtle research. While limitations and room for further improvement remain, the addition of our platform and general approach to the marine turtle genetic toolbox opens the door to a diversity of rapid, cost-efficient genotyping applications. These data can be comparable across laboratories, geographical regions, and timescales, which can be particularly important in such highly mobile species that can migrate across entire ocean basins and necessitate international collaboration for effective conservation (Shamblin *et al*. 2014). Though our specific selected regions for targeted enrichment will not be suitable for all populations or research questions, our study also demonstrates how initial RAD-Sequencing can be used to develop a Rapture platform suited to specific research needs. Additionally, these target regions can be adapted to other genotyping platforms that may be better suited to meet some research needs but require prior knowledge of genomic variants, e.g., GT-Seq that may have improved performance on lower quality and concentrations samples (Campbell *et al*. 2015) or microhaplotypes that may provide increased power for relationship inference (Baetscher *et al*. 2017).

Our results highlight several key strengths of this platform in meeting the diverse needs of marine turtle genotyping applications. First, researchers often need to analyze few or many samples at few or many loci, depending on study goals. Our data demonstrate that samples can be combined and effectively genotyped at the same loci with moderate sequencing coverage using partial capture reactions. This not only facilitates cost-effective, time-efficient analysis of large sample sets, but also combining samples for different projects. For example, researchers working on large nesting beaches often have many samples to analyze at the end of the season (Shamblin *et al*. 2017), while those genotyping samples from fisheries bycaught animals or some foraging population assessment projects may have smaller sample sets collected intermittently over the year. In the latter case, it has been particularly problematic to determine how to move from manual analysis with traditional markers to next-generation sequencing approaches where much of the reduced cost and time efficiency is related to multiplexing and high-throughput processing. While genotyping high priority single samples that need to be analyzed in near real-time may still pose a challenge, the flexibility of the Rapture platform offers options to combine library preparation and sequencing across projects and species, or to create a libraries with fewer samples and reduce total sequencing depth (e.g., through the use of a lower output instrument such as an Illumina MiSeq, or coordinating with other researchers to use different library barcodes and share sequencing lanes). Additionally, we designed a custom MYBaits enrichment kit with ∼2000 targets to satisfy the needs of a variety of study types, but this can be adapted to include fewer or more loci. For example, researchers interested in basic population structure and individual assignment may wish to design kits with a subset of only several hundred informative targets, increasing the per locus depth of coverage in each sample. Finally, the ability to repeatedly capture the same genomic regions facilitates studies conducted over broader time periods (e.g., examining trends across many nesting seasons or even generations) or spatial scales (e.g., collaborating labs can generate and share data between foraging and nesting grounds).

Despite these exciting opportunities, our data also clearly show that our current Rapture platform has some limitations that are relevant to situations frequently encountered in wildlife genetics studies. First, although we were able to effectively perform high on-target sequencing and genotyping for samples across tissue types, DNA extraction methods, species, and other co-factors, a portion of our test samples failed to sequence well. Though no clear patterns emerged with sample age or molecular weight thresholds, it is likely that highly degraded or contaminated samples (e.g., due to natural conditions, collection and storage methods) were more likely to fail. While this problem is often easily circumvented in controlled experimental settings, in many conservation applications these issues can be unavoidable, such as working with museum collections or opportunistic sampling of animals that have had substantial exposure to natural elements postmortem. However, we emphasize that many samples in our study that exhibited evidence of some degradation were successful, including those that fall into these sub-optimal categories (e.g., stranded and bycaught animals). Our results support the initial findings of Ali *et al*. (2016) that this new RAD protocol is more robust than previous RAD methods for partially degraded samples, but there may be a point beyond which it is not a suitable approach. However, it may be possible to generate comparable genotype data for these samples at a subset of informative Rapture loci with highly-multiplexed PCR based methods such as GT-Seq (Campbell *et al*. 2015) that amplify short DNA fragments and thus be more robust to sample degradation. Secondly, we observed a substantial proportion of sequenced fragments that were PCR clones, and this was correlated with initial sample DNA concentration. The latter observed effect may be a product of the increased influence of measurement and pipetting error at low concentrations, which could be targeted for improvement in a future protocol adaptation. However, since PCR clones are in effect wasted sequences, in practice this currently means that it is less cost effective to sequence samples with low initial DNA concentrations, and that calculations of required sequencing to attain a targeted depth of coverage must take these factors into account. Although sequencing costs are likely to continue to decrease such that genotyping can still be achieved despite this loss, future efforts to reduce clonality would improve the efficiency and cost of this approach. Finally, although costs and technological accessibility have vastly improved in recent years, access to the equipment and financial resources to conduct genetic studies is far from universally available. This makes continued collaboration essential to advancing our understanding of marine turtles, with researchers with access to such resources working to increase capacity elsewhere, such as through visiting scientist training partnerships and creation of shared genetic databases. Particularly given the influence that bioinformatics parameters (e.g., filtering criteria, assembly methodology, genotyping thresholds) can have on results (O’Leary *et al*. 2018), it is imperative for researchers to include metadata and analysis details to ensure robust and comparable data across laboratories and over time.

We present results of conducting SNP discovery independently for each species and within a representative leatherback population to demonstrate that substantial variation exists within our targeted regions to meet a variety of study goals, but also to highlight the importance of appropriate test data and analyses parameter thresholds to avoid ascertainment bias (i.e., discerning informative SNPs appropriate for a given study goal; Lachance & Tishkoff 2013). For example, intra-population questions can require variable SNPs within a target population, which may not be identified in broader analysis including many populations depending on filtering thresholds and sample sizes (Andrews *et al*. 2018). One advantage to the flexible Rapture platform is that researchers can generate data for many genomic regions and then hone in on informative SNPs to genotype without *a priori* knowledge and the need to develop different markers tailored to each study goal, which can be cost and time prohibitive. However, as discussed previously, if desired, researchers can also use preliminary RAD or Rapture data with a representative test dataset to identify the most informative markers for their study and design new MYBaits kit or GT-Seq primers to focus exclusively on those targets.

Principal components and admixture proportion analyses identified clear separation of all species examined and our tree depicting relationships among species was in general agreement with previous research (Duchene *et al*. 2012; Naro-Maciel *et al*. 2008). It is important to note that these studies were focused on resolving phylogenetic relationships among all marine turtle species, and thus the methods employed were much more in-depth than our analyses; additionally, we were not able to include any flatback turtle samples in our study. Thus, clarifying any discrepancies or further confirmation using our genome-wide markers would require additional studies. However, for the purpose of our primary study goals, since species were randomized across and within RAD libraries and we observed low number of sequences assigned to blank wells, our results show that sequences can be assigned correctly to individuals using this highly-multiplexed approach and our analyses criteria. Cross-species targeted enrichment may not be as effective in other taxa with high genomic diversity or for studies that require tens to hundreds of thousands of SNPs, and researchers working with other species may wish to omit targets from our panels that only yielded coverage in green or leatherback turtles.

We identified several hybrids, in agreement with preliminary evaluation of these samples with three nuclear loci and the mitochondrial control region (Dodge *et al*. 2006), though additional analyses with larger sample sizes from contributing species at the same locations would further validate these findings and provide insight into the prevalence of hybridization in these populations. Hybridization and complex introgression patterns have been previously documented, primarily in southeast Atlantic populations (Reis *et al*. 2010; Vilaça *et al*. 2012), but the frequency of such events elsewhere and the relative hybrid fitness is largely unknown. Given recent concern that increasingly skewed female-biased sex ratios due to climate change (Jensen *et al*. 2018) and other anthropogenic pressures (Gaos *et al*. 2018) could cause interspecies mating events to become more prevalent and further destabilize populations, additional research is needed to better understand these processes and monitor changes over time; our Rapture platform offers an additional tool for such studies

Our exploratory green turtle analyses determined that our platform can also successfully amplify targeted regions within species across broad geographic locations and identify informative SNPs for stock structure, population assignment and other management applications. A recent study of green turtle global phylogeography using mtDNA control region sequences identified eleven divergent lineages that each encompass a few to many genetically differentiated distinct management units (MUs) with more recent shared ancestry but deemed to be demographically independent (Jensen *et al*. in press). This comprehensive study builds on previous work within regions documenting restricted gene flow attributed to female natal philopatry and generally little genetic differentiation among nesting beaches within 500km (reviewed in Jensen *et al*. 2013; Jensen *et al*. in press; Komoroske *et al*. 2017). While instrumental for our understanding of green turtle evolutionary history and contemporary stock structure patterns, there is a clear need to complement this work with studies employing nuclear markers to identify the roles of male-mediated gene flow and higher marker resolution. With additional refinement of the SNPs identified here specifically to meet these goals (e.g., narrower filtering criteria to remove any biases due to physical linkage or inconsistent coverage), these markers will serve as a valuable resource for such studies over large spatial and temporal scales, further advancing our understanding of green turtle population connectivity, MU designation, and human impacts.

Finally, comparisons of genetic variation among populations and species can be informative for a variety of conservation relevant research, such as understanding how genetic diversity may differ among healthy, recovering, and declining populations (Lozier 2014). While our current sample set was not designed to address these questions specifically, the ability to consistently amplify over a thousand regions across the genome for all marine turtles, enables our platform can be effectively employed for such research goals within or across species. For example, we found that Pacific leatherbacks exhibited the lowest levels of nucleotide diversity relative to all other groups evaluated, including the (Atlantic) Brazilian nesting stock. While further robust analysis is needed to confirm this preliminary finding, this could be related to the continued decline of Pacific leatherback populations in contrast to Atlantic populations.

In conclusion, our Rapture platform provides a tool that is complementary to existing traditional genetic markers as well as other emerging genomic techniques suited to address a broad diversity of research questions in marine turtle ecology, evolution and conservation (e.g., transcriptome, other reduced representation, and whole genome sequencing to study adaptive variation and genome-phenome linkages). Though some limitations still hinder widespread adoption of these techniques, such as cost and well-assembled and annotated genomic resources, as technologies continue to advance we anticipate continued application and creative adaptations to meet the challenging needs of conservation researchers. If realized, this could generate capacity for large-scale initiatives such as the creation of global genetic databases akin to those that have begun emerging recently for other taxa (e.g., Deck *et al*. 2017). This would not only expand the scope of research questions that can be investigated, but also provide traditionally resource-limited marine turtle programs with the ability to incorporate genetic information in their research and monitoring efforts. Such endeavors will inevitably present many new challenges, but the successes of analogous initiatives such as the State of the World’s Sea Turtles (SWOT) and the Atlantic-Mediterranean Loggerhead Genetics (LGWG; Shamblin *et al*. 2014) working groups among others have demonstrated the power of such global collaborative efforts to answer the major outstanding research questions in these wide-ranging, complex megafauna.

## Acknowledgements

We would like to thank members of the Marine Mammal and Turtle Division for assistance in the laboratory, logistics and project design suggestions, especially A. Frey, P. Morin, V. Pease, E. LaCasella, A. Lanci, G. Serra-Valente and S. Roden. Funding was provided by the National Oceanic and Atmospheric Administration, Southwest Fisheries Science Center, the National Research Council of the National Academies of Science, Engineering and Medicine to LMK, through a Lenfest Ocean Program grant to KRS (the views expressed are those of the authors and do not necessarily reflect the views of the Lenfest Ocean Program or The Pew Charitable Trusts), and The Sea Turtle Census Initiative at The Ocean Foundation. We would like to thank T. Summers, D. Graff, A. Tagarino, N. Fitzsimmons, A. Gaos, M. Liles, K. Dodge, H. Guzman, N. Marcovaldi, J. Thome, B. Bowen, J. Mortimer, E. Harrison and G. Balazs, Centro Tamar/ICMbio and Projeto Tamar/Fundação Pro Tamar, The Sea Turtle Conservancy, American Samoa Department of Marine and Wildlife Resources, CNMI Division of Fish and Wildlife & Department of Lands & Natural Resources and NOAA-Fisheries Pacific Islands Science Center, for sample contributions.

## Data Accessibility

Data analyses scripts, documentation and Rapture platform probe sequences are available at https://github.com/lkomoro/Marine_Turtle_Rapture_Methods. Illumina raw reads for Trial 2 hardshell turtles are deposited in NCBI Sequence Read Archive (Bioproject PRJNA487648).

## Author Contributions

LMK, MM, SO, MPJ, KRS and PHD contributed to the conceptual design of the project. LMK, MM and SO conducted laboratory, marker design, and data analyses. LMK, MPJ, KRS and PHD assessed data interpretation for green turtles, and LMK and PHD wrote the manuscript.

